# Recovery of genomes from metagenomes via a dereplication, aggregation, and scoring strategy

**DOI:** 10.1101/107789

**Authors:** Christian M. K. Sieber, Alexander J. Probst, Allison Sharrar, Brian C. Thomas, Matthias Hess, Susannah G. Tringe, Jillian F. Banfield

## Abstract

Microbial communities are critical to ecosystem function. A key objective of metagenomic studies is to analyse organism-specific metabolic pathways and reconstruct community interaction networks. This requires accurate assignment of assembled genome fragments to genomes. Existing binning methods often fail to reconstruct a reasonable number of genomes and report many bins of low quality and completeness. Furthermore, the performance of existing algorithms varies between samples and biotopes. Here, we present a dereplication, aggregation and scoring strategy, DAS Tool, that combines the strengths of a flexible set of established binning algorithms. DAS Tool applied to a constructed community generated more accurate bins than any automated method. Further, when applied to environmental and host-associated samples of different complexity, DAS Tool recovered substantially more near-complete genomes, including novel lineages, than any single binning method alone. The ability to reconstruct many near-complete genomes from metagenomics data will greatly advance genome-centric analyses of ecosystems.

## Introduction

Genome-resolved metagenomics targets the reconstruction of genomes from environmental shotgun DNA sequence data. Based on the genome sequence, metabolic pathways of individual organisms can be inferred and their lifestyle in the microbial community can be predicted. The challenge of recovering genomes from complex mixtures of sequence fragments is comparable to that of assembling jigsaw puzzles from a mixture of many puzzles without knowing how many puzzles are present and what they look like. Not surprisingly, powerful bioinformatics methods are required to achieve the desired outcome.

Existing binning methods use features derived from sequence composition, sequence abundance or taxonomy inferred from reference databases. Early methods primarily made use of shared GC content and coverage (sequence depth) to cluster together the fragments belonging to specific genomes^1^. As the complexity of ecosystems targeted for analysis increased, additional methods became essential. Teeling *et al*. proposed the use of sequence compositional information, primarily tetranucleotide frequencies, as a binning input^2^. This approach made use of genome characteristics established through study of the genomes of isolated organisms^3^. Sequence compositional analysis (tetranucleotide and other k-mer composition and codon usage data) was implemented within emergent self-organizing maps (ESOMs) to successfully extract genomes from metagenomes^4^. The ESOM-based approach has been widely used to recover draft genomes from many different environments^5,6^, but it rarely works well if the dataset contains fragments from a large number of different organisms (as is typical of soil and sediment). The ESOM method is also somewhat subjective, as the cluster boundaries are user-defined.

A major advance in binning methods came with the realization that the pattern of organism abundances across a sample series was a binning signature^7,8^. This approach assumes that contigs of one organism have a similar abundance (as measured by mapped read counts) in one sample and that the representation of all contigs from a genome should change in the same way across a sample series.

Phylogenetic profile information was of minimal use early in the metagenomics era because the number of reference microbial genomes was very small (a few dozen genomes). However, the phylogenetic signal of a contig that derives from sequence similarity is now a useful constraint for binning of data from some samples, and it continues to grow in utility as the number of reference genome sequences (from isolates, single cells and genomes from metagenomes) increases.

Current state of the art automated binning tools combine sequence abundance and composition into one model^9,10^ and some of them additionally use marker genes from a reference database^11^. The quality and completeness estimation of the output of automated binning tools is essential. CheckM, for example, tests for a set of single copy marker genes to determine the completeness of bins and give an estimate of the amount of contamination of a bin^12^.

Existing binning tools are based on broadly accepted features and clustering algorithms. Additionally, the tools were all benchmarked using datasets analysed in their respective publications. In fact, most binning methods were demonstrated using relatively simple communities (e.g., premature infant gut datasets of Sharon *et al*.^7^). However, the value of bins generated when these methods are applied to other samples is uncertain, and bins generated when different tools are applied to a new dataset may differ significantly in completeness and contamination. Furthermore, some genomes may be exclusively predicted by just one tool. Here, we tested the performance of a set of well-established binning methods by applying them to data from a group of ecosystems that varies dramatically in complexity. We found that no single tool or approach performed well on all ecosystems. Furthermore, many incomplete bins and multi-genome mega-bins were predicted. The different performance of binning tools and the fact that different tools reconstruct different genomes with varying levels of completeness motivated the development of a strategy that integrates the results of predictions of multiple binning algorithms. Probst *et al*. combined the results of three binning methods in a comparative approach with additional manual curation and increased the total number of reconstructed near-complete genomes from a subsurface aquifer environment over that obtained using just one method^13^. However, because different binning predictions are based on the same assembly of contigs, predicted bin overlap was extensive and the determination of an optimal consensus draft genome set was not trivial. These findings motivated the development of the dereplication, aggregation and scoring tool (DAS Tool). DAS Tool is an automated method that integrates a flexible number of binning algorithms to calculate an optimized, non-redundant set of bins from a single assembly. We show that this approach generates a larger number of high quality genomes than achieved using any single tool.

## Results

### Development of an integrative binning approach

The DAS Tool approach to solve the binning problem is to integrate predictions from multiple established binning tools. The number and type of binning tools is flexible. Candidate bins are generated independently when all binning tools are applied to the same assembly. DAS Tool then uses a consensus approach to select a single set of non-redundant, high quality bins. The approach relies on two major components: (1) A scoring function that estimates the quality and completeness of the bins. The score is based on the presence, absence, and number of duplicated single copy genes in a bin, making it possible to compare predictions from different methods and to select an optimal bin-set. (2) An algorithm that extracts a non-redundant and optimized bin set from multiple binning predictions based on the scoring function (Figure 1). DAS Tool selects the highest scoring bin from the candidate set and assigns it to the final bin set. The scaffolds of that bin are then removed from all other bins in the candidate set. The scores of all of the candidate bins from which scaffolds were removed are then updated. The selection procedure iterates until all remaining candidate bins have values that do not exceed a pre-defined minimum bin quality threshold (calculated based on the scoring function). Nevertheless, we advise that the user examine each of the final bins to identify potential contamination based on erroneous phylogenetic affiliation and to remove sequences from phage/virus (based on gene content).

**Figure 1.**
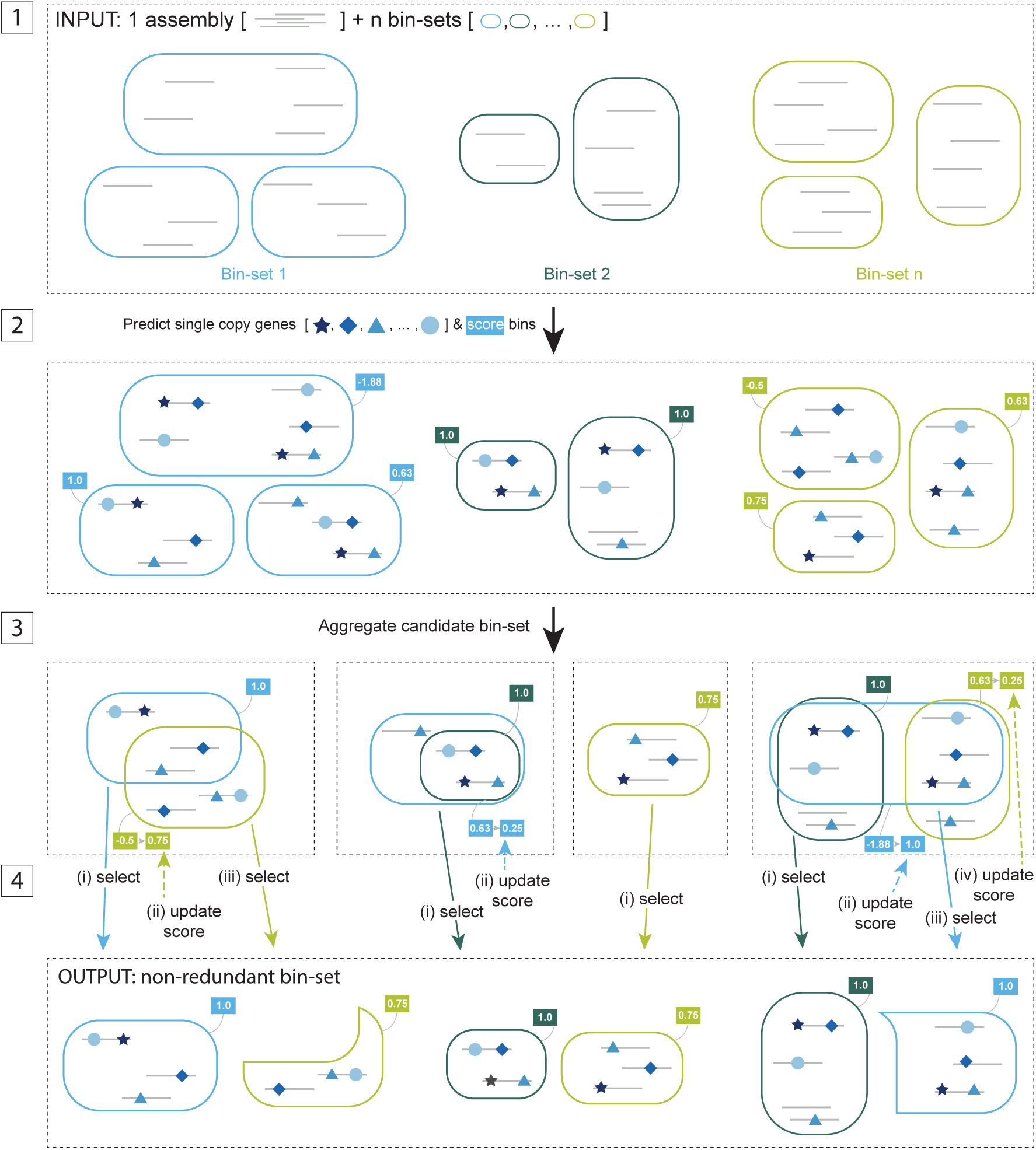
Overview of the DAS Tool algorithm. Step 1: Input of DAS Tool is scaffolds of one assembly (grey lines) and a variable number of bin-sets from different binning predictions (rounded rectangles of same color). Step 2: Single copy genes (blue shapes) on scaffolds are predicted and scores (blue and green boxes) are assigned to bins. Step 3: Aggregation of redundant candidate bin-set from all binning predictions. Step 4: Iterative selection of high scoring bins and updating of scores of remaining partial candidate bins. Output comprises non-redundant set of high scoring bins from different input predictions.

## DAS Tool applied to a synthetic community comprised of a mixture of isolates

To validate the DAS Tool algorithm, we applied it to data from a synthetic microbial community that was constructed by mixing together DNA of 22 bacteria (including different species from the same genus) and 3 archaea^14^. We predicted bins using five binning tools (ABAWACA 1.07 (https://github.com/CK7/abawaca), CONCOCT^9^, MaxBin 2^11^, MetaBAT^10^ and tetranucleotide ESOMs^4^) and combined the result using DAS Tool. In addition, we manually binned the genomes using ggKbase binning tools^15^ (ggkbase.berkeley.edu) that make use only of GC, coverage and taxonomic profile.

To determine how well the reconstructed bins represent the reference genomes, we calculated F_1_ scores, which is the harmonic mean of precision and recall. In addition we estimated the completeness of bins based on marker genes using CheckM^12^. Bins were only considered to be of high (>90% complete) or draft (70% - 90% complete) quality if they had less than 5% contamination due to the presence of multiple genes expected to be in single copy. Many predicted bins with high F_1_ scores also were classified as high quality by CheckM, based on completeness and contamination (Figure 2 a, b). With a F_1_ score above 0.9 for all 25 reconstructed bins, ggKbase performed best on the synthetic community. However, this result is only generalizable to a few other systems (see below for examples) because its success is based on the clear phylogenetic signal from reference genomes in public databases. Of the other predictions, the bins reported by DAS Tool show the highest accuracy in terms of F_1_ scores. DAS Tool reports 12 bins with an F_1_ score above 0.99, followed by tetranucleotide ESOMs and MaxBin 2 with 9 bins. Similarly, DAS Tool reports 20 bins with an F_1_ score above 0.95 followed by tetranucleotide ESOMs with 18 bins (Figure 2 a, Supplementary Table 1).

**Figure 2.**
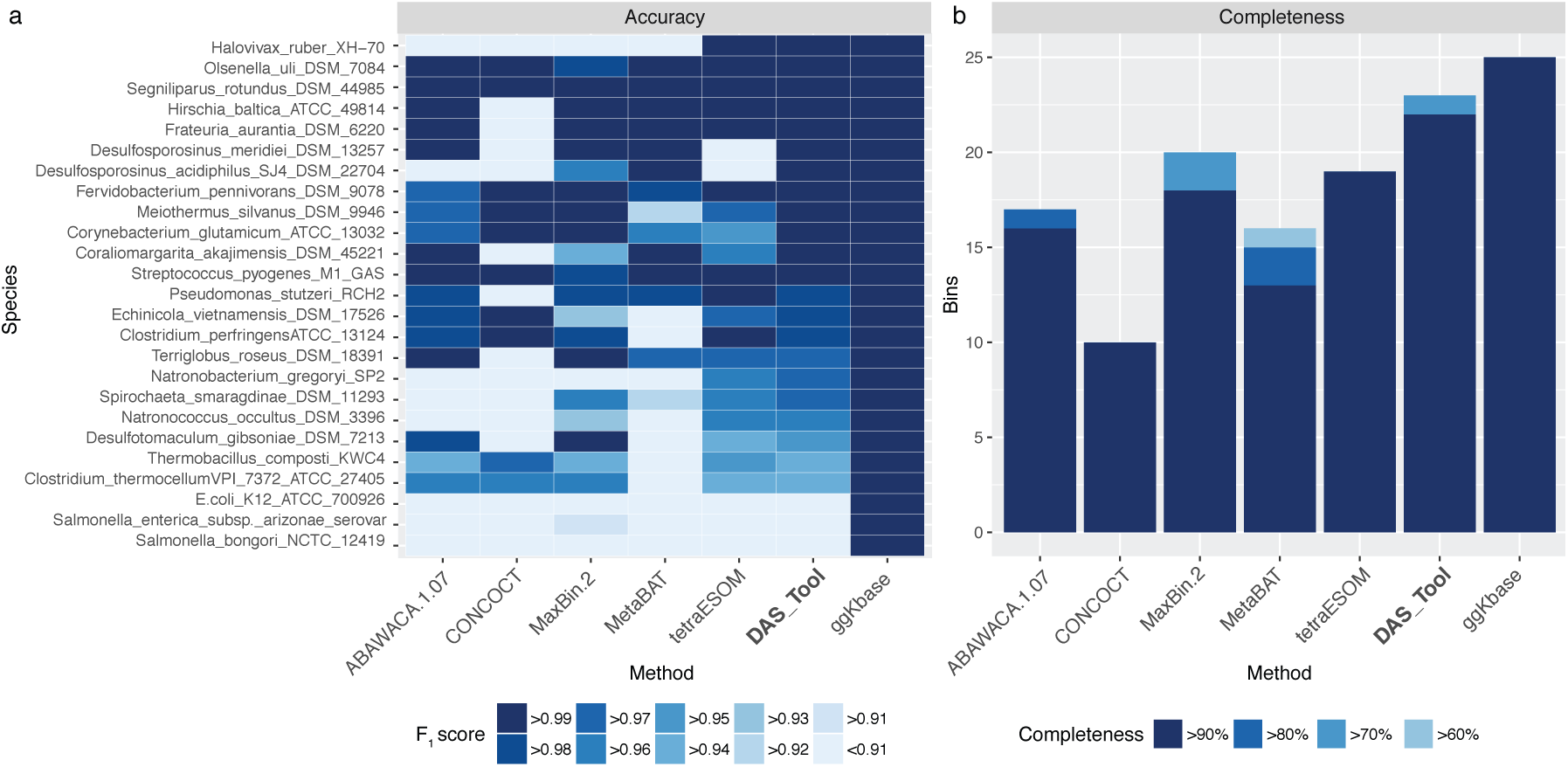
Reconstructed genomes from a synthetic mock community consisting of 25 isolate genomes. (a) Accuracy of reconstructed genomes per method based on F_1_ score. (b) Number of reconstructed high quality genomes with low contamination (< 5%) according to marker gene based completeness estimation^12^.

DAS Tool not only has the highest accuracy in terms of the F_1_ score metric but also reports the highest number of near-complete genomes with low contamination. Only the manual binning approach using ggKbase was able to reconstruct all 25 genomes.

## Application of DAS Tool to environmental metagenomic data

Probst *et al*.^13^ generated a highly curated set of genome bins from metagenomic data from a high CO_2_ cold-water geyser that were ideal for evaluation of the DAS Tool algorithm. The data comprise two assemblies of sequences from samples collected sequentially on 3.0-µm and 0.2-µm filters and a set of 3.0-µm filtrates from subsurface fluids collected at a single time point. The published bins were generated by a comparative approach of three methods followed by manual curation of the results^13^. We used CheckM to generate quality estimates for the published bins that can be compared to quality estimates for all binning methods, including DAS Tool.

We compared the results of the three independent binning predictions from Probst *et al*. (ABAWACA 1.0, tetranucleotide ESOMs, differential abundance ESOMs), as well as those from ABAWACA 1.07, CONCOCT, MetaBAT, and MaxBin 2 to results achieved using DAS Tool. DAS Tool was applied using either a combination of three or seven different binning algorithms (Figure 3, Supplementary Table 2).

**Figure 3.**
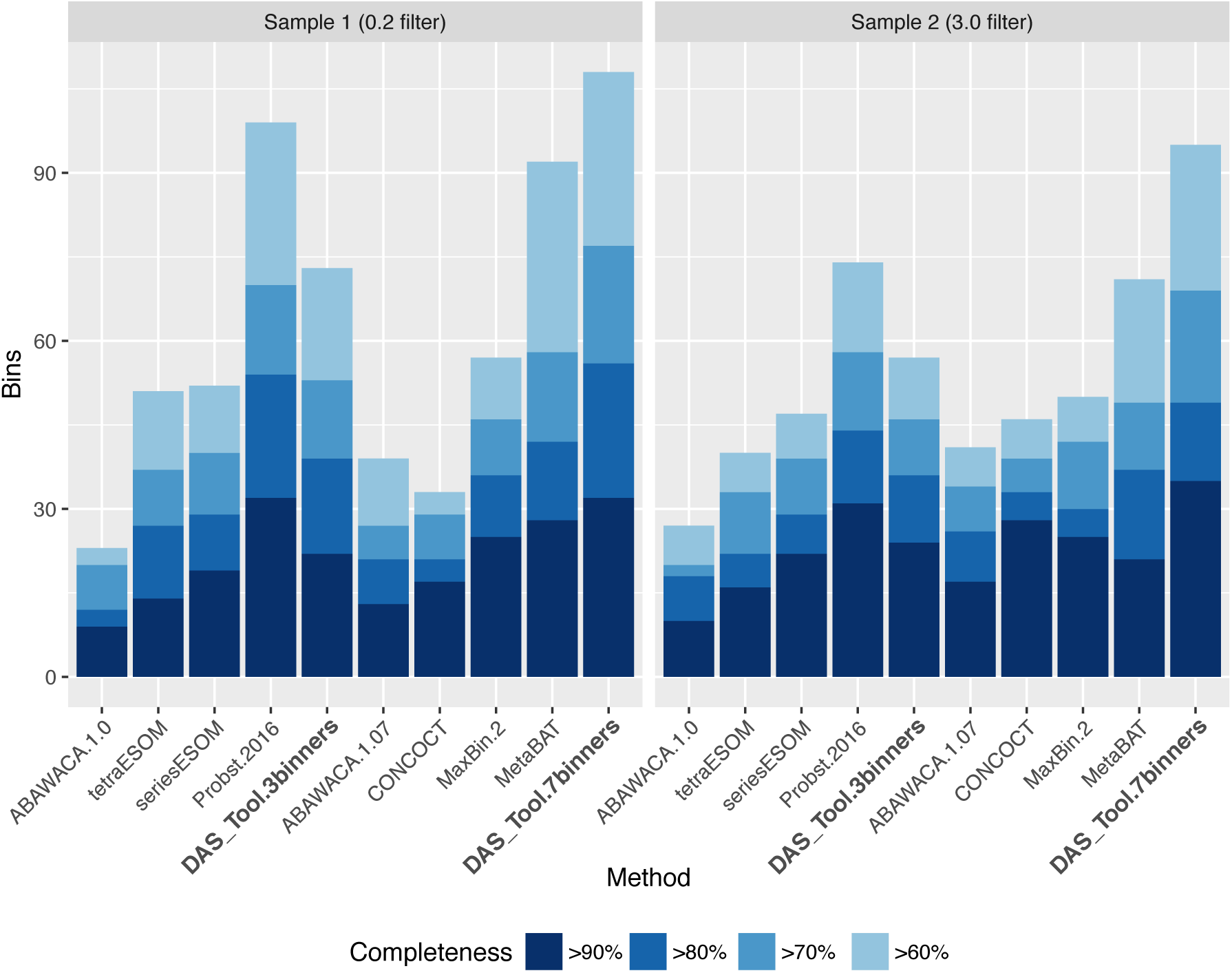
Reconstructed genomes from Crystal Geyser, a high CO_2_ cold water geyser. Number of high quality genomes with low contamination (<5%) from metagenomic assemblies of two samples. Probst.2016 represents the combination by Probst et al., 2016^13^ of ABAWACA.1, tetraESOM and seriesESOM and a final manual curation step. DAS_Tool.3binners uses the same three predictions as input. DAS_Tool.7binners additionally uses ABAWACA.2, CONCOCT, MaxBin.2 and MetaBat.

Although DAS Tool with three binning algorithms reported more near-complete and draft genomes than the three methods alone, it returned less genomes than in the curated set by Probst et al. (Figure 3, Supplementary Table 2). However, when we included seven binning tools in DAS Tool (adding ABAWACA 1.07, CONCOCT, MaxBin 2 and MetaBAT), the reported number of near-complete genomes was the same for the 0.2-µm sample (32 genomes) and even higher for the 3.0-µm sample (DAS Tool: 35, Probst: 31). For both samples a larger number of draft genomes was reconstructed than was achieved previously^13^ (Figure 3, Supplementary Table 2). The number of draft genomes increased slightly when allowing more contamination per bin (Supplementary Figure 1).

## Combination of bins using DAS Tool improves genome count from metagenomic data with different levels of complexity

In order to evaluate the performance of DAS Tool on samples of different complexity, we applied it to shotgun metagenomic data of lower, medium and high complexity from human microbiomes^16^, natural oil seeps^17,18^, and soil (see Data Availability). We binned all samples separately using ABAWACA 1.07, CONCOCT, MaxBin 2, MetaBAT and tetranucleotide ESOMs. All predictions were combined using DAS Tool and CheckM was used to estimate the quality of the resulting bins. In addition, we used ggKbase binning tools to analyse the human gut data. This was appropriate, given colonization of the human gut by genomically well-characterized bacteria. ggKbase tools were not used in the other analyses because they do not perform well in systems with high phylogenetic novelty (data not shown).

Summing up the number of bins of each quality level that were generated for the three ecosystems, DAS Tool reported the highest number of near-complete and draft bins in all cases (Figure 4).

**Figure 4.**
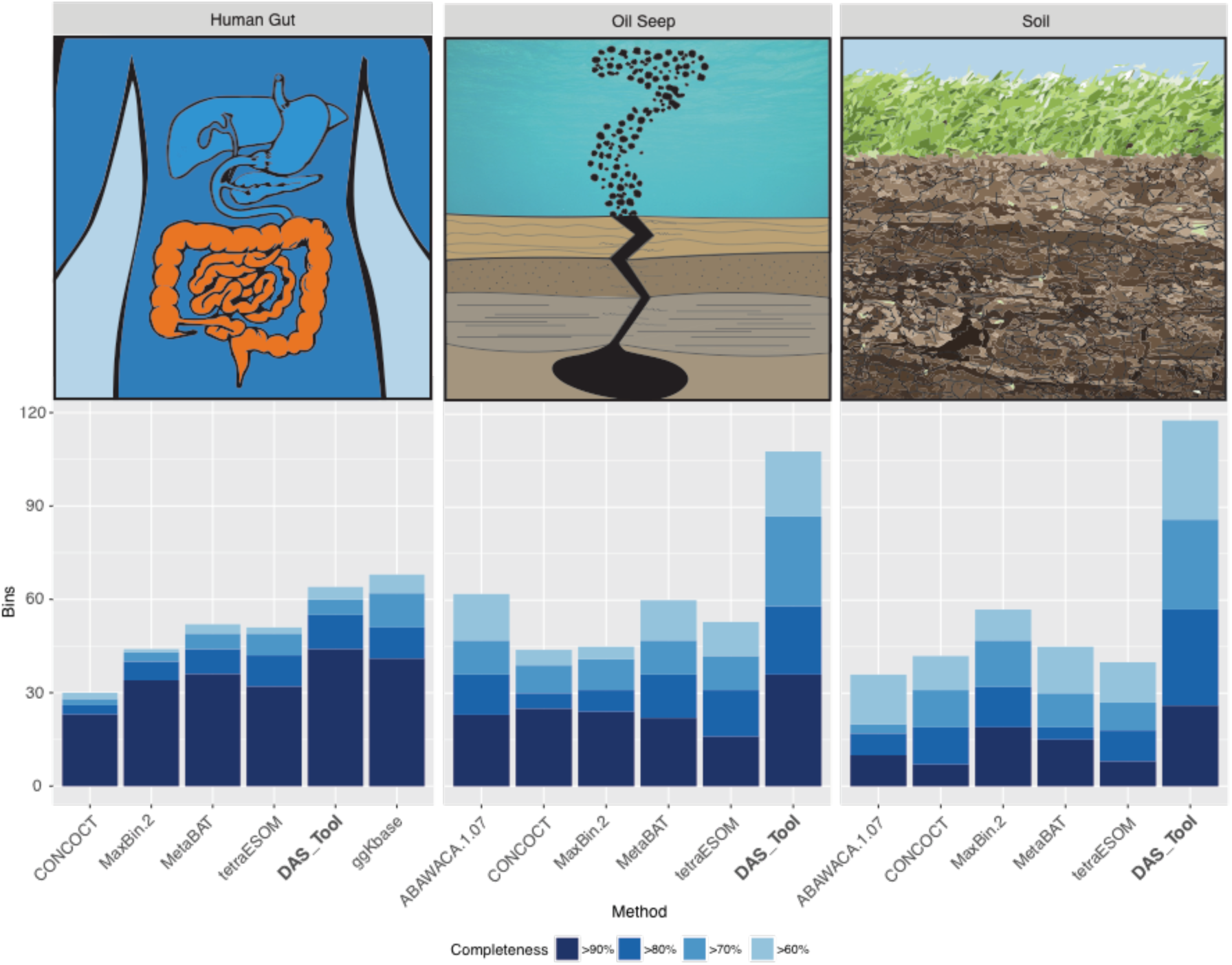
Number of high quality genomes with low contamination (< 5%) from metagenomic assemblies of samples from three ecosystems representing a range of complexity. Samples were collected from adult human gut (1 fecal sample), oil seeps (5 samples), and hillslope soil and underlying weathered shale (6 samples). Samples were assembled and binned separately. Reconstructed genomes were summed up per ecosystem. For sample-by-sample results, see **Supplementary Figure 2**.

Interestingly, the performance of the single binning tools that were used as input for DAS Tool (which excludes ggKbase) differed between ecosystems and none of them was the clear “winner”. In the case of bins generated for the lower complexity human gut samples using single binning tools, ggKbase followed by MetaBAT generated the largest number of near-complete genomes. For the medium complexity oil seeps, ABAWACA 1.07 and MetaBAT produced the most draft-quality genomes while CONCOCT produced slightly more high quality bins. For high complexity soil data MaxBin 2 reported the most draft and near-complete genomes. Compared to the best performing method, DAS Tool reports 1.2, 1.4, and 1.4 times more near-complete genomes and 1.2, 1.9 and 1.8 times more draft quality genomes for human gut, oil seeps and soil samples, respectively (Figure 4).

We also examined the performance of the various binning approaches sample by sample. DAS Tool reported either the most or the same number of near-complete genomes with low contamination for 11 out of 13 samples (higher: 7/13, equal: 4/13). It generated up to 1.5 times more bins than the best performing single binning method. For draft quality bins, DAS Tool generated the largest number of bins for 12 out of 13 samples, and up to twice as many draft quality bins than the best performing single binning method (Supplementary Figure2). The number of reconstructed genomes per sample increases when considering genomes with a higher amount of contamination. In 10 out of 13 samples (higher: 9/13, equal: 1/13) DAS Tool reports more or the same number of genomes with more than 70% completeness and less than 15% contamination (Supplementary Figure 3, Supplementary Table 3).

## Genome analysis reveals novel lineage with hydrocarbon degradation potential

Binning of metagenomic data from Santa Barbara oil seep samples revealed three genomes, whose 16S rRNA sequences lacked closely related sequences in the SILVA database^19^ (78.8%, 79.4% and 87.4% identity). The estimated completeness of these newly reconstructed genomes ranges from 95.6% to 89.6% (Supplementary Table 4).

In a phylogenetic tree based on 16 concatenated ribosomal proteins, the three genomes cluster as a monophyletic group with one TA06 and two WOR-3 genomes (Supplementary Figure 1 a). The JGI_Cruoil_03_Bacteria_38_101 forms a cluster together with the TA06 lineage at a patristic distance of 1.2977 but is more distant to the two WOR-3 (patristic distances: 1.5531 and 1.5258, respectively). In contrast, the two lineages JGI_Cruoil_03_Bacteria_44_89 and JGI_Cruoil_03_Bacteria_51_56 share greater similarity with the two WOR-3 at a minimal patristic distance of 1.3350 and 1.0582, respectively and have a greater distance to the TA06 (patristic distance: 1.4328 and 1.4673, respectively).

For comparison, the patristic distance between representatives of closely related phyla in the same tree was between 1.0282 and 1.2110 (Firmicute *Thermincola sp.* JR versus the Chloroflexus *C. aurantiacus* j-10-fl and Melainabacteria *Obscuribacter phosphatis* versus the Cyanobacteria *Leptolyngbya sp.* PCC 7104).

Given that both distances are smaller than the distances of TA06 and WOR-3 to our reconstructed genomes JGI_Cruoil_03_Bacteria_38_101 and JGI_Cruoil_03_Bacteria_44_89 as well as the distance of JGI_Cruoil_03_Bacteria_38_101 to JGI_Cruoil_03_Bacteria_44_89 (patristic distance: 1.5164) we conclude that these two new genomes may be representatives of two new phylum-level lineages. The third novel genome JGI_Cruoil_03_Bacteria_51_56 is closer to the WOR-3 at a patristic distance of 1.0582 and is likely part of the WOR-3 candidate division.

Interestingly, the 16S rRNA gene sequences of all three of our newly reconstructed novel genomes group with some sequences classified as TA06 and one sequence classified as a WS3 (the other WS3 sequences form a lineage sibling to Zixibacteria) (Supplementary Figure 1 b). Except for one TA06 (Candidate_division_TA06_bacterium_32_111), the corresponding TA06 and WS3 genomes place distant from our genomes on the concatenated ribosomal protein tree. Thus, some of the 16S rRNA gene sequences of these publicly available genomes may be misclassified or misbinned (a common problem with 16S rRNA gene binning, especially if the gene is in multi-copy and the scaffolds are short). Regardless, it is clear that our genomes are highly distinct from any other genomes in public databases.

Pathway analysis reveals genes encoding for hydrocarbon degradation enzymes, including aldehyde dehydrogenase, which is present in all three genomes. Additionally, alcohol dehydrogenase, aldehyde ferredoxin oxidoreductase and methanol dehydrogenase are present in JGI_Cruoil_03_Bacteria_44_89, the genome with highest estimated completeness, suggesting pathways for degradation of alkanes and methanol (Supplementary Table 5).

## Genomes from soil

From six soil samples, we reconstructed 82 minimally contaminated (<5%) draft genomes (>70% completeness), 24 of which were high quality draft genomes (>90% completeness) (Supplementary Figure 2). Two of the high quality genomes were well-assembled (a Gemmatimonadetes genome consisting of 11 scaffolds and a Bacteroidetes genome on 14 scaffolds), with estimated completeness above 97% and contamination below 3.3%.

It has been shown recently that some Gemmatimonadetes are able to consume methanol using a pyrrolo-quinoline quinone (PQQ)-dependent methanol dehydrogenase (MDH) and to convert the resulting formaldehyde using the tetrahydromethanopterin (THMPT) and tetrahydrofolate (THF)-linked formaldehyde oxidation pathways^20^. Likewise, we were able to find a PQQ-MDH and two key enzymes of the THF pathway (methenyltetrahydrofolate cyclohydrolase, methylenetetrahydrofolate dehydrogenase) in the high quality Gemmatimonadetes genome bin but could not find any enzymes belonging to the THMPT pathway. Additionally, we found genes for carbon fixation, fermentation, nitrogen assimilation, complex carbon degradation, and sulfur metabolism. Similarly, the Bacteroidetes genome encodes enzymes for carbon fixation, fermentation, and nitrogen assimilation but by contrast has no genes for methane metabolism, complex carbon degradation, or sulfur metabolism (Supplementary Table 5).

## Discussion

We tested a group of currently available, published metagenomics binning algorithms to evaluate how well they performed when applied to samples of a wide range of complexity. Consistent with previous work showing that use of data series signals can significantly improve binning outcomes^7,8^, the single binning algorithms that used these signals (CONCOCT, MaxBin, MetaBAT, ABAWACA) performed better than composition-based tools (tetra-ESOM) on most samples. However, it is notable that each of these was variably effective across the different system types, and even among different samples from the same ecosystem, and no single binning algorithm was consistently the most effective. Interestingly, for very simple communities that include organisms that are closely related to genomically-characterized species (e.g. the synthetic community), the manual combination of phylogeny, GC, coverage and single copy gene inventory produces good binning outcomes; however, this is not the case for more complex datasets.

In contrast, DAS Tool, the new consensus binning strategy presented here, almost always extracted considerably more genomes from complex metagenomes than any of the single binning tools alone. While DAS Tool did not outperform manual bin combination and curation when using the same starting set of bins from three single binning approaches, adding four additional binning algorithms resulted in equal or more near-complete bins than the published manually curated results. This finding underlines the advantage of including more binning methods in DAS Tool. It is important to note, even tools that generate only a small number of high quality bins can significantly improve the result of DAS Tool because other tools sometimes miss these bins.

It is not uncommon for the research community to question the quality of genomes reconstructed from metagenomes. Imperfect bins are a challenge for all studies that attempt to genomically resolve complex ecosystems. However, if they can be obtained, the value of high quality draft genomes is enormous. Different single algorithm methods not only generate different numbers of bins but the genome content can differ slightly. This variable performance can be evaluated by using strategies such as DAS Tool. In picking the best bins from each binning tool, DAS Tool is able to equalize performance variations of single binning tools and thus increase the total number of near-complete genomes recovered. Because it uses a single copy gene based scoring function it is able to distinguish between high and low quality bins and by using an appropriate score cutoff it can filter out low quality bins and control the number of megabins.

Despite improvements in assembling and binning methods, reconstructing genomes from soil metagenomics data is still seen as challenging. With the help of DAS Tool we were able to extract dozens of high quality genomes from soil, including some near complete genomes. Furthermore, in re-analysing public data from off shore oil seep sediments we identified and genomically characterized organisms of a novel lineage that is likely involved in hydrocarbon degradation.

In conclusion, DAS Tool can integrate manual binning methods such as emergent self-organizing maps (ESOMs) and can incorporate the results of any contig-based binning algorithm. Thus, it is highly scalable and can make use of binning tools developed in the future.

## Methods

### Implementation

DAS Tool is implemented in R^21^. Besides R-base functions, we used the R-packages doMC^23^ to implement multicore functionality, data.table^24^ for efficient data access and storage and ggplot2^25^ to visualize results. DAS Tool is available under https://github.com

### Scoring function

To estimate the quality and completeness of predicted bins we set up a scoring function (Equation 1). The function calculates a bin score based on the frequency of 51 bacterial or 38 archaeal reference single copy genes (rSCG). The first term of the function represents the fraction of single copy genes (SCGs) present and accounts for the completeness of the genome. It is the number of unique single copy genes per bin (uSCG) divided by the number of reference SCGs (rSCG). The second term accounts for contamination and decreases the score in case of duplicated SCGs (dSCG). It is calculated by the ratio of the unique number of duplicated SCGs (dSCG) divided by the total number of unique SCG (uSCG) in a bin. The third term is a penalty for megabins and is the total number of extra single copy genes divided by the number of reference genes. It is calculated by the difference of the total number of predicted SCGs (ΣSCG) and the number of unique SCGs per bin divided by the number of reference SCGs. Both penalty terms are accompanied by weighting factors. For this study we set b=1.5 and c=1 to favour low contaminated bins.

For each bin scores using the bacterial and archaeal reference gene set are calculated and the greater of the two scores is reported as bin score.

Equation 1 Scoring function

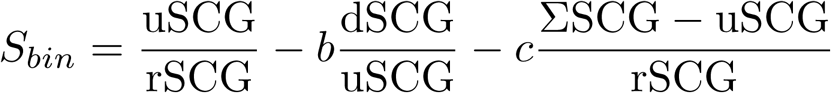

### Marker gene prediction

Genes in the assembly are predicted using prodigal^26^ with the meta option and the ‘-m’ flag for preventing gene models to be built over ambiguous nucleotides. Single copy marker genes (SCGs) are determined in using databases of bacterial^27^ and archaeal SCGs^13^ as a seed to select candidates of single copy genes from the metagenomes using USEARCH^28^ (e-value 1e-2). The candidates were then searched^28^ against the entire database (e-value 1e-5) and called present if the query spanned at least 50% of the alignment with the best hit in the database.

Although all results shown in this manuscript are based on USEARCH^28^, DAS Tool can also make use of the open source tools DIAMOND^29^ and BLAST^30^ to predict single copy genes. Scripts for SCG prediction are available under https://github.com/AJProbst/sngl_cp_gn.

### Selection algorithm

In the first step, a redundant candidate bin set is created, which consists of all predicted bins of the input binning methods. The quality of all bins in the candidate set is estimated using the SCG-based scoring function (Equation 1). After that in an iterative procedure a non-redundant bin set is selected (Figure 1). First the highest scoring bin is extracted out of the candidate set. If two or more bins have the same score, the bin with a higher scaffold N50 value is chosen. If the N50 value is also equal, the larger bin in terms of nucleotide sequence is selected. After removing the bin out of the set, also all contigs that belong to this bin are removed out of other bins. Because this step influences the composition of other bins, the scoring function is applied again on all altered bins. The iteration continues as long as selected bins are above a pre-defined score threshold (default: 0.1) or until all bins in the candidate set are selected.

### Data availability

The reads of human gut samples (SRA accession: SRR3496379)^16^, Crystal geyser samples (BioProjects PRJNA229517 and PRJNA297582) and the synthetic community (SRA accession: SRX1836716)^14^ were obtained from NCBI. Reads of the oil seep data (Gold Analysis Project IDs: Ga0004151, Ga0004152, Ga0004153, Ga0005105, Ga0005106)^17,18^ and soil samples (Gold Analysis Project IDs: Ga0007435, Ga0007436, Ga0007437, Ga0007438, Ga0007439, Ga0007440) were downloaded from JGI portal pages (https://img.jgi.doe.gov/cgi-bin/m/main.cgi. Assemblies were downloaded from ggKbase for the human gut samples (http://ggkbase.berkeley.edu/LEY3/organisms) and from IMG for the oil seep samples (Gold Study ID: Gs0090292). Genomes from oil seep and soil samples, which were analysed in this study, are available on ggKbase (http://ggkbase.berkeley.edu/dastool) and NCBI ([TBD]).

### Assembly and mapping

The reads of the synthetic community and soil samples were quality filtered by SICKLE (Version 1.21, https://github.com/najoshi/sickle, default parameters) and assembled using IBDA_UD^31^. All samples were assembled separately. Read mapping for all samples was done using Bowtie 2^32^.

### Binning

For generating input bin sets for DAS Tool, we applied the automated binning tools ABAWACA 1.07 (https://github.com/CK7/abawaca), CONCOCT^9^, MaxBin 2^11^ and MetaBAT^10^. We also calculated tetranucleotide ESOMs^4^ and selected clusters manually using Databionic ESOM Tools^33^. Additionally, we manually binned the low complexity synthetic community and the human gut microbiome data based on GC, coverage and taxonomic profile using ggKbase tools^15^ (http://ggkbase.berkeley.edu). Bins predicted by ggKbase were only used for comparison purpose and not used as input for DAS Tool. ABAWACA 1.07 returned no results on the human gut data due to the lack of differential coverage information. The bins of ABAWACA 1.0, tetranucleotide ESOMs and differential-abundance ESOMs for the Crystal Geyser data was obtained from Probst et al.^13^.

### Binning evaluation

We used a synthetic community of 26 genomes^14^ for evaluating the accuracy of binning predictions. The genome of *Nocardiopsis* was not considered for this analysis as low sequence coverage (0.54% based on mapping^14^) did not allow its reconstruction by the assembler. The assembly was mapped on the remaining 25 reference genomes using NUCmer^34^ and used to calculate F_1_ scores, which is the harmonic mean of precision and recall. Besides that, we estimated marker gene based completeness of bins using the lineage workflow of CheckM^12^.

### Genome curation and annotation

Assemblies of submitted genomes were error corrected using re_assemble_errors.py (https://github.com/christophertbrown/fix_assembly_errors). Gene prediction was performed with the same settings used for marker gene prediction in DAS Tool (prodigal^26^ in meta mode and ‘-p’ flag). Functional predictions were made using the ggKbase annotation pipeline, which uses USEARCH^28^ to search predicted ORFs against Kegg^35^, UniRef100^36^ and UniProt^37^.

### Phylogenetic tree

The ribosomal protein tree is based on concatenated alignments of the amino acid sequences of 16 ribosomal proteins (ribosomal protein L2, S3, L3, L4, L5, L6P-L9E, L15, L16-L10E, S8, L14, L18, L22, L24, S10, S19 and S17). Alignments were created for each protein using MUSCLE^38^ and trimmed manually. After concatenation columns with more than 95% gaps were removed. We calculated the phylogenetic tree using the maximum likelihood algorithm RAxML^39^ on the CIPRES web server^40^ in choosing the LG (PROTCATLG) evolutionary model and autoMRE to automatically determine the number of bootstraps. 16S rRNA gene sequences were aligned using SSU-align^41^, trimmed and submitted to the CIPRES web server^40^. We used RAxML^39^ and the GTRGAMMA model and determined the number of bootstraps using autoMRE. Patristic distances, which are the sum of branch lengths between two taxa in a phylogenetic tree, were calculated using the cophenetic.phylo function of the ape R-package^42^.

### Code availability

DAS Tool is available under https://github.com/cmks/DAS_Tool (v1.0 used in this analysis).

## Acknowledgements

We thank Itai Sharon for support for the new ABAWACA version, Karthik Anantharaman, Edward Kirton, and Adam Rivers for inspiring discussions, Bill Andreopoulos for technical support, Spencer Diamond and Matt Olm for beta testing.

This work was supported by the Emerging Technologies Opportunity Program of the U.S. Department of Energy Joint Genome Institute, a DOE Office of Science User Facility, supported under Contract No. DE-AC02-05CH11231. Support was provided by the DOE grant DOE-SC10010566 and NIH grant 5R01AI092531. Work by A.J.P. was supported by the DFG grant PR1603/1-1.

### Author contributions

C.M.K.S. designed and implemented the DAS Tool algorithm. A.J.P. and B.C.T. provided scripts for the DAS Tool upstream analysis. C.M.K.S., A.J.P., and A.S. performed data analyses. M.H. provided Santa Barbara oil seep data. B.C.T. and J.F.B. provided ggKbase pipeline annotation and phylogenetic assignments. J.F.B. binned the synthetic and the human gut community using ggKbase. C.M.K.S. and J.F.B. wrote the paper with contributions from S.G.T., A.J.P. and M.H. All authors reviewed the results and approved the manuscript.

### Supplementary Material

#### 1. Supplementary figures

**Supplementary Figure 1.**
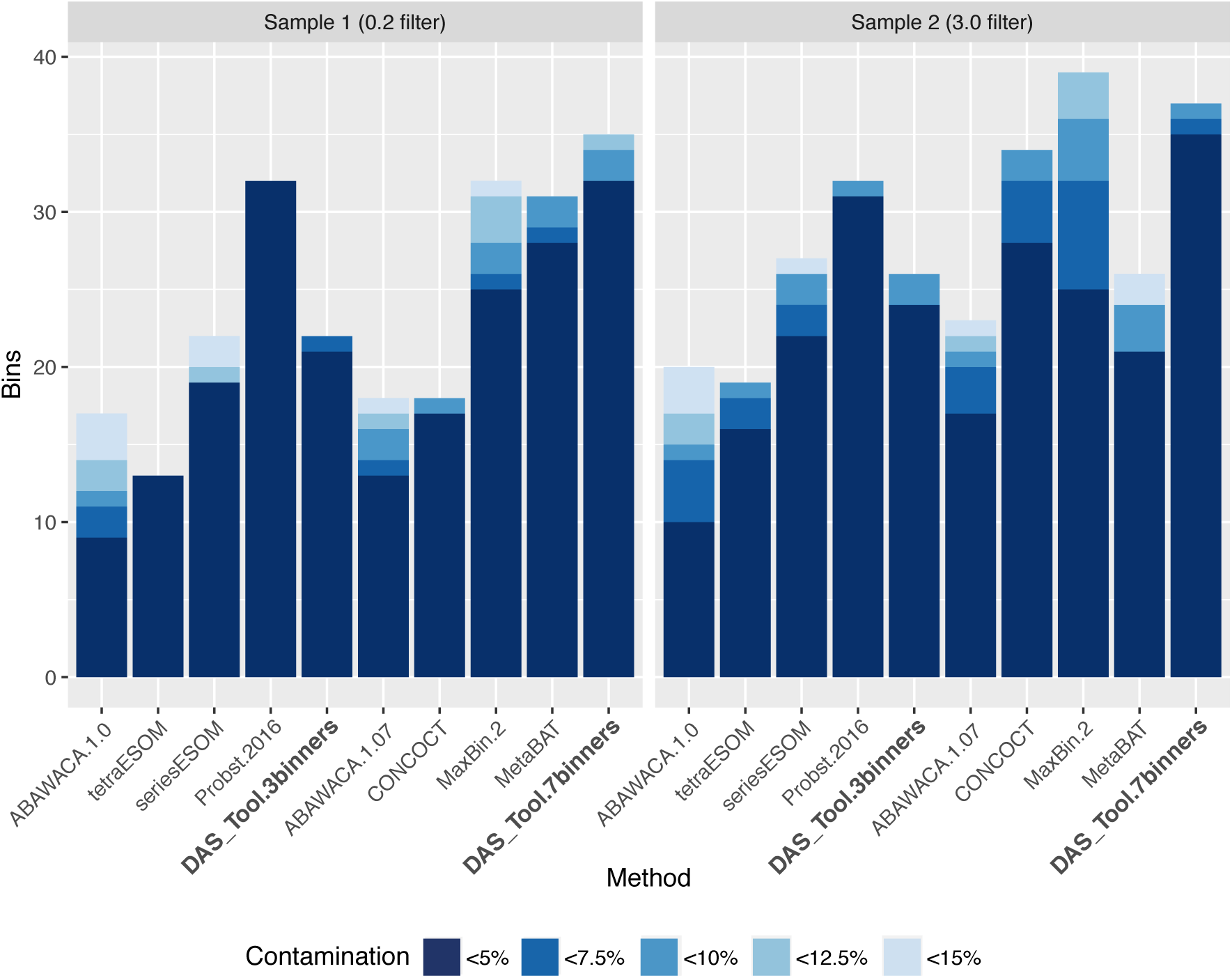
Number of draft genomes with at least 70% completeness and less than 15% contamination for two real metagenomic assemblies from Crystal Geyser, a high CO_2_ cold water geyser.

**Supplementary Figure 2.**
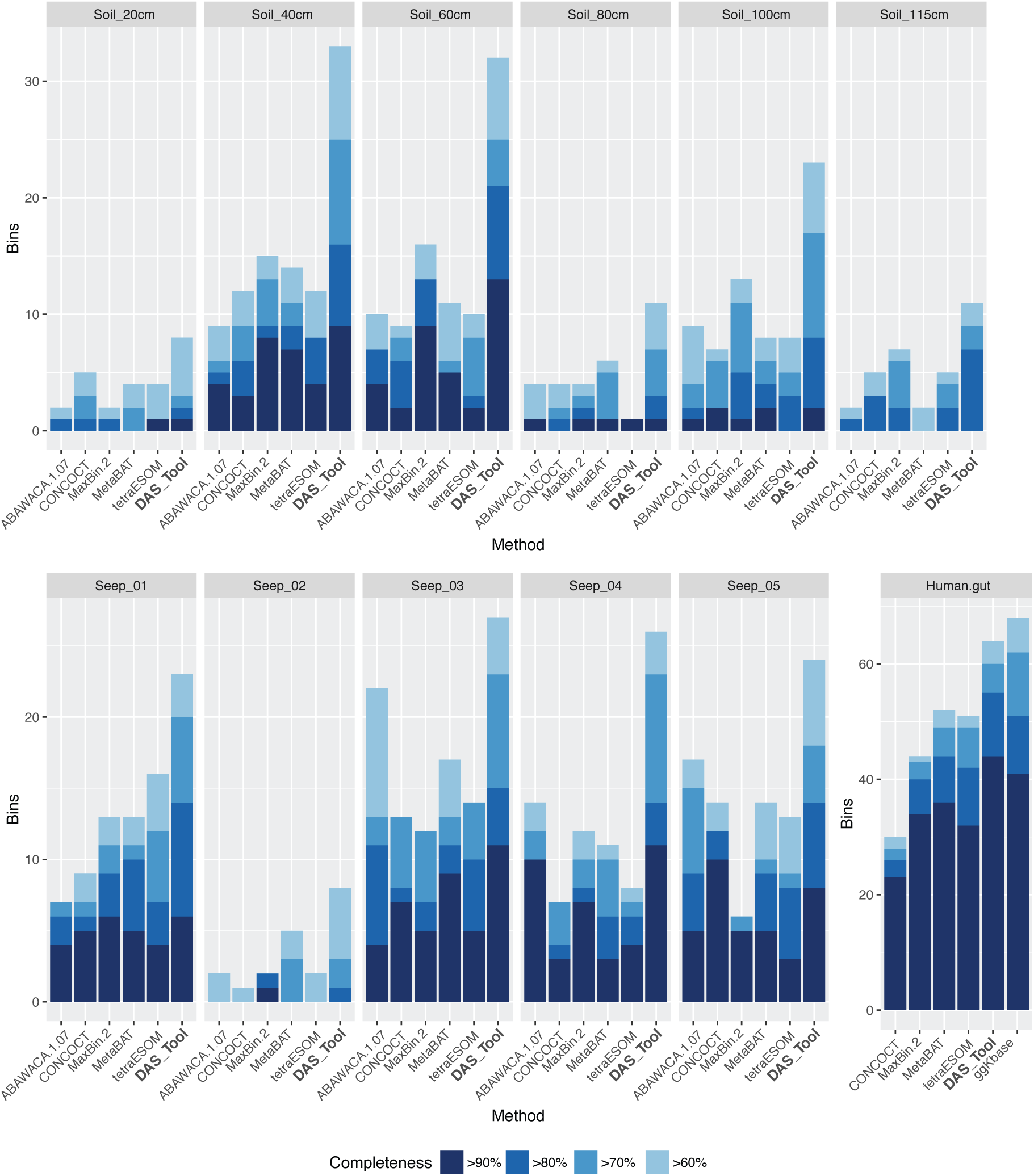
Number of high quality genomes with low contamination (<5%) for twelve real metagenomic assemblies representing a range of complexity. Samples were collected from adult human gut, oil seeps and hillslope soil.

**Supplementary Figure 3.**
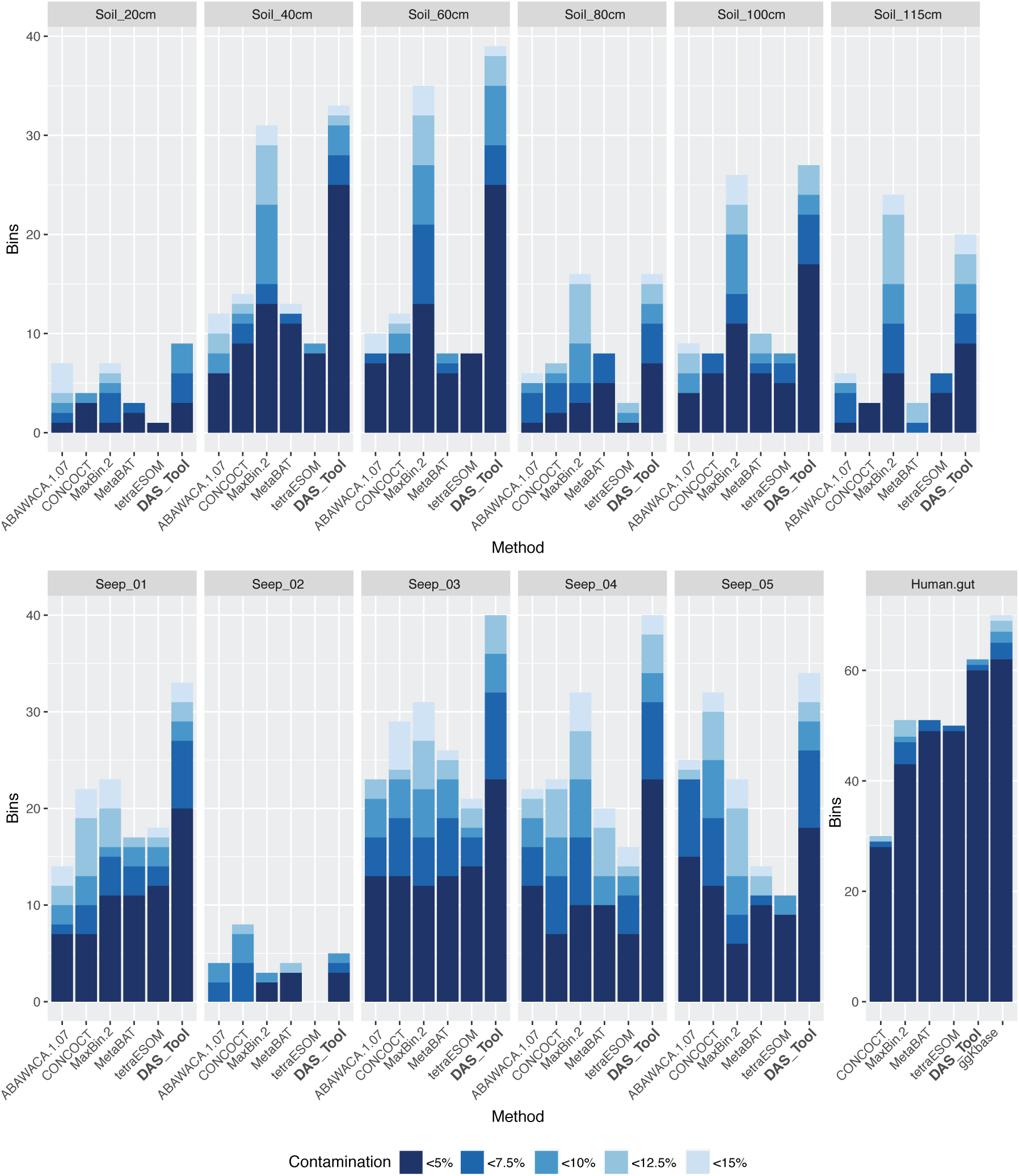
Number of draft genomes with at least 70% completeness and less than 15% contamination for twelve real metagenomic assemblies representing a range of complexity. Samples were collected from adult human gut, oil seeps and hillslope soil.

**Supplementary Figure 4.**
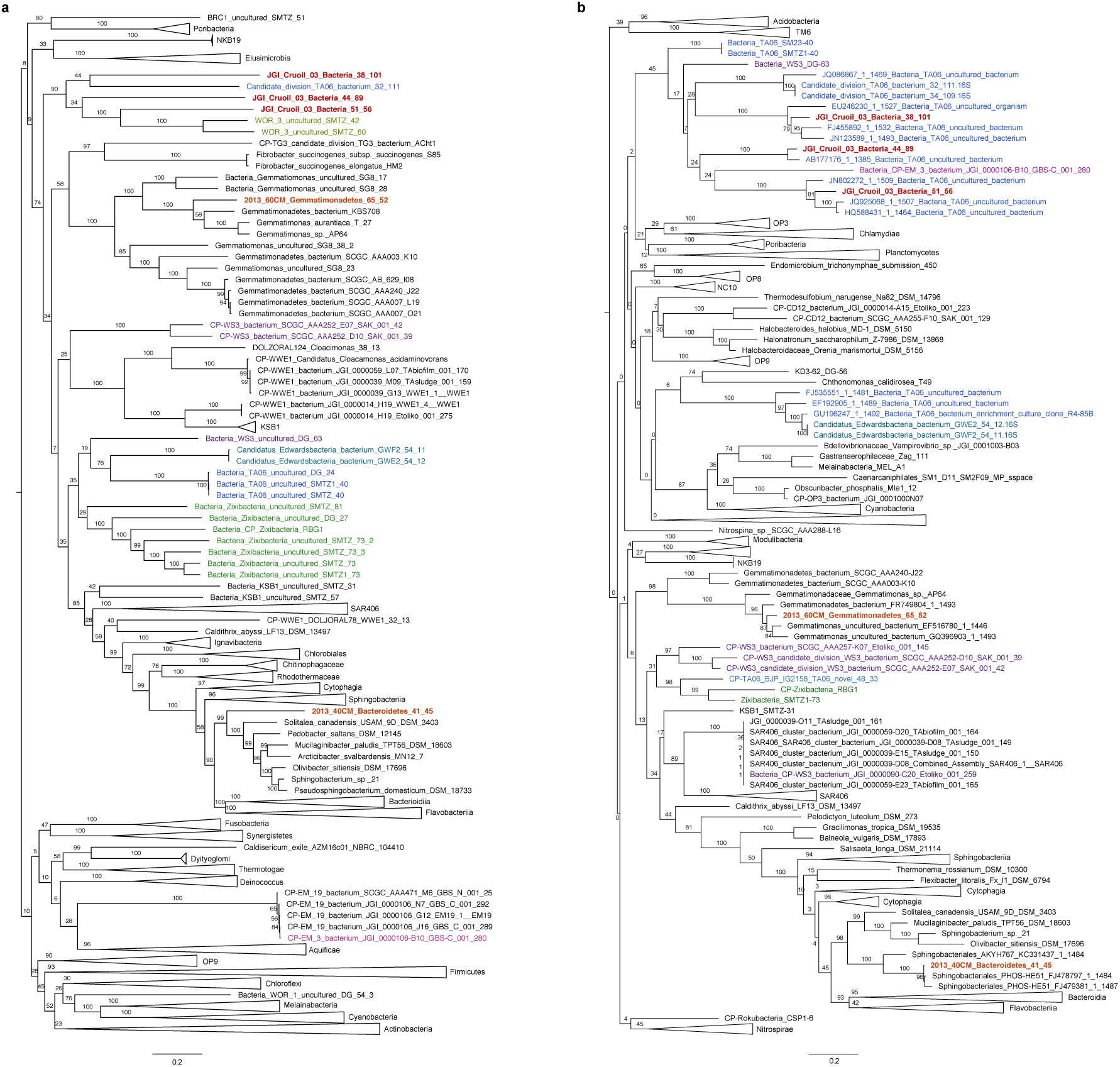
Phylogenetic trees based on 16 concatenated ribosomal protein sequences (a) and based on 16S rRNA sequence showing reconstructed genomes from oil seeps (red) and soil metagenomes (orange). Reference genomes include TA06 (blue), Edwardsbacteria (emerald), WOR-3 (olive), WS-3 (purple), EM-3 (magenta) and Zixibacteria (green).

#### 2. Supplementary tables

**Supplementary Table 1.** Accuracy of reconstructed genomes from a synthetic mock community consisting of 25 isolate genomes based on F_1_ score.

**Supplementary Table 2.** Reconstructed genomes from Crystal Geyser, a high CO2 cold water geyser. Number of high quality genomes with low contamination (< 5%) from metagenomic assemblies of two samples. Probst.2016 represents the combination by Probst et al., 2016 of ABAWACA.1, tetraESOM and seriesESOM and a final manual curation step. DAS_Tool.3binners uses the same three predictions as input. DAS_Tool.7binners additionally uses ABAWACA.2, CONCOCT, MaxBin.2 and MetaBat.

**Supplementary Table 3.** Number of high quality genomes with low contamination (< 5%) from metagenomic assemblies of samples from three ecosystems representing a range of complexity. Samples were collected from adult human gut (1 fecal sample), oil seeps (5 samples), and hillslope soil and underlying weathered shale (6 samples). Samples were assembled and binned separately. Reconstructed genomes were summed up per ecosystem.

**Supplementary Table 4.** Genome quality estimates (CheckM) and 16S sequence similarities(SILVA) of reconstructed genomes from oil seeps.

**Supplementary Table 5.** Predicted key enzymes of metabolic pathways of five reconstructed genomes from oil seep and soil metagenomes.

